# Maize Stem Buckling Failure is Dominated by Morphological Factors

**DOI:** 10.1101/833863

**Authors:** Christopher J Stubbs, Ryan Larson, Douglas D Cook

## Abstract

The maize (Zea mays) stem is a biological structure that must both balance biotic and structural load bearing duties. These competing requirements are particularly relevant in the design of new bioenergy crops. With the right balance between structural and biological activities, it may be possible to design crops that are high-yielding *and* have digestible biomass. But increased stem digestibility is typically associated with a lower structural strength and higher propensity for lodging. This study investigates the hypothesis that geometric factors are much more influential in determining structural strength than tissue properties. To study these influences, both physical and *in silico* experiments were used. First, maize stems were tested in three-point bending. Specimen-specific finite element models were created based on x-ray computed tomography scans. Models were validated by comparison with in vitro data. As hypothesized, geometry was found to have a much stronger influence on structural stability than material properties. This information reinforces the notion that deficiencies in tissue strength could be offset by manipulation of stalk morphology, thus allowing the creation of stalks with are both resilient and digestible.

**Highlight:** This study utilized physical and *in silico* experiments to confirm that geometric parameters are far more influential in determining stalk strength than mechanical tissue stiffnesses.

## Introduction

As plant stems grow, they must balance both their biological and physical functions (Badel *et al.*, 2015). These functions are balanced based on their perceived need, leading to plant stem systems that grow more or less depending on their experienced external abiotic stimuli *(Moulia et al., 2015)*. As such, if plant stems do not properly acclimate to their wind-loading environment, wind-induced bending can cause failure (Niklas and Spatz, 2012). This problem is compounded in bioenergy maize varieties, where stem digestibility often results in lower strength stems that are more susceptible to win-induced bending failure. The three primary failure modes of grain stems are compressive tissue failure at the outer fiber, transverse buckling, and longitudinal splitting (Schulgasser and Witztum, 1992; Wegst and Ashby, 2007; Robertson *et al.*, 2015*a*). Mature dried maize stems can exhibit all three failure modes, but compressive tissue failure and transverse buckling are the most common (Robertson *et al.*, 2015*a*).

It is well known that a reduction in lignin causes crop tissues to be more easily digested for biofuel production (Simmons *et al.*, 2010), but can also cause several wholistic agricultural deficiencies (Pedersen *et al.*, 2005). In a previous paper, we reported that mechanical stresses within the maize stalk were “much more sensitive to changes in physical dimensions than they were to changes in the material properties” (Von Forell *et al.*, 2015). This suggested that changes to stalk morphology could be used counteract the reduction in tissue properties, thus potentially enabling crops that were both digestible and structurally robust. But that paper relied upon a relatively limited set of computational models without direct empirical support and acknowledged that further experimentation would be required to fully explore the mechanisms of stalk strength.

Multi-scale modeling can be extremely useful in developing a mechanistic understanding of the causes and modes of failure of structural systems (Júnior *et al.*, 2011). A key challenge in the multi-scale modeling approach with respect to structural systems is that the relationship among parameters across the array of modeling length scales is not immediately obvious (Hufner and Accorsi, 2009). Furthermore, the impact of parameters on the overall structural behavior of the system is often obfuscated by the covariance of multiple parameters. Because of this, it can be difficult or impossible to isolate the impact of individual parameters on the behavior of the stalk through purely experimental/statistical approaches. Computational modeling and engineering theory allow us to gain a deeper mechanistic insight into the failure modes of stalks while also informing research between modeling length scales.

The purpose of this study was to test the ideas put forth in our 2015 paper by using both physical and *in silico* experiments to more clearly determine the influence of the geometry and material properties on the structural robustness of maize stems. Another previous study from our research group identified relationships between morphological features of maize stems and their bending strength (Robertson *et al.*, 2017). This study looks to expand on those two previous studies with more detailed specimen-specific finite element models to broaden our understanding of the effects of geometry and material properties on maize stem strength.

## Materials and Methods

### Overview of the process

This study involved three-point bending tests of maize stalk specimens and corresponding specimen-specific *in silico* finite element experiments. Maize specimens approximately one meter in length with low moisture content were tested in three-point bending according to a previously described testing protocol (Robertson *et al.*, 2015*b*). Prior to testing, x-ray computed tomography (CT) scans were performed. Specimen-specific finite element models were created from CT scan data. Models were validated against experimental data and then used to perform flexural stiffness and buckling analyses. Material properties were varied to assess the influence of various parameters on the flexural stiffness of the models. The influence of geometry was evaluated using a statistical analyses of the three-point bending tests in conjunction with geometric data from CT scans.

### An Overlapping Experimental/In Silico Approach

A previous study from our lab describes the scanning and testing of 980 maize stalk specimens (Robertson *et al.*, 2017). The experiments and data collected in that study were used as a starting point for this study. Results from previous studies indicated that the major diameter, minor diameter, and rind thickness were the primary factors that influenced both flexural stiffness and stalk strength (Von Forell *et al.*, 2015; Robertson *et al.*, 2017). Furthermore, the values of these factors in the internodal region of the stalk were most predictive of flexural stiffness as well as overall strength (Robertson *et al.*, 2017). The major diameter, minor diameter, and rind thickness in the internodal regions were therefore used as geometric parameters in this study.

Geometric and material sensitivities were assessed through the use of *in silico* experiments. This approach was used because the measurement of material properties is very labor-intensive and typically requires dissection of the specimen. But material properties are easily modified *in silico*, and the opportunity to vary each material property independently significantly reduces the sample size required to obtain meaningful insights. Specimen-specific *in silico* experiments were based upon individual specimens from within the sample of stalks tested experimentally. Models were validated by comparing the predicted vs. measured values of flexural stiffness.

However, because the geometry of the maize stalk is complex and unique to each specimen, the sensitivity of maize stalks to geometric factors was assessed using experimental data. Regression analyses were used to determine the sensitivity of flexural stiffness and bending strength to geometric factors derived from CT data of the corresponding stalks.

### Model development

In our three-point bending tests, buckling failure always occurred near the central node. As a result, computational expense was minimized by modeling only the stalk section in the vicinity of the failure region. The length of the finite element continuum model was 100 mm, which adequately captured the buckling failure modes while also minimizing Saint-Venant effects (Sandorff, 1980; Stubbs *et al.*, 2018). Shear and bending loads were applied to the end faces of each finite element model. These loads were calculated from the loading configuration of the 3-point bending experiments; see Figure 1. With loads applied in this manner, the loading anvil was be assumed to be immobile. As a result, the point of contact between the anvil and the stalk was fixed in all 6 degrees of freedom. Although this constraint was required for model stability, it also imposed rotational constraints. The resultant moments at the point of contact were found to be negligible, thus indicating that this boundary condition did not adversely influence the validity of the model.

**Figure 1.**
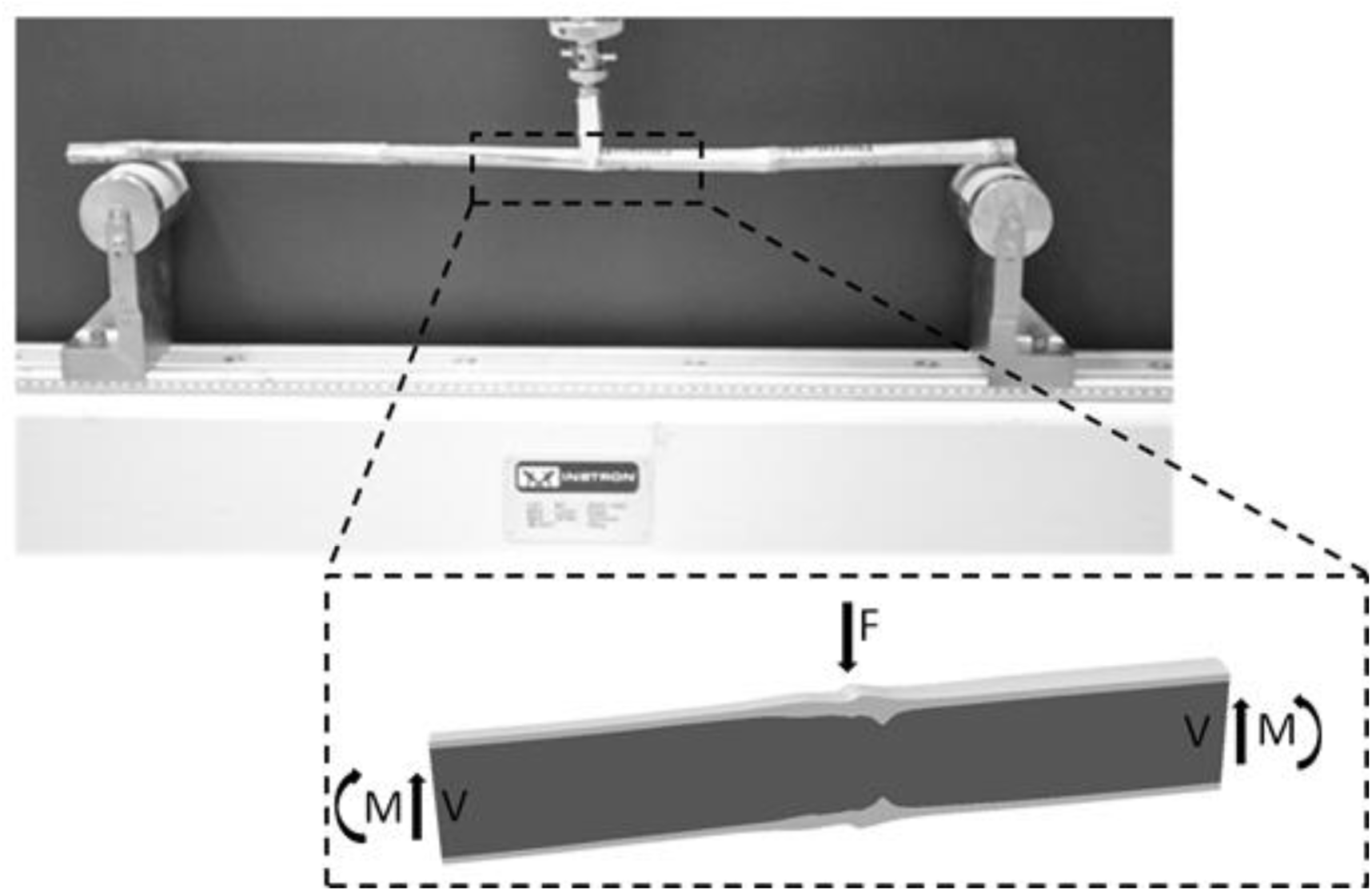
The 100 mm long specimen-specific solid continuum models were created.

All finite element models were developed in Abaqus/CAE 2016. The geometry was meshed using 10-noded quadratic tetrahedral elements (Matthews *et al.*, 2000; Hibbitt *et al.*, 2016; Simulia, 2016). The flexural stiffness analyses used a linear, full-Newton direct solver. The linear buckling analyses used a subspace eigensolver. A more detailed description of the finite-element modeling process is available in the supplementary information, including the development of the validation models.

### Material properties

The material of both the rind and pith was modeled as transversely isotropic. Using the force-displacement response of the specimen from the bending experiments, a flexural stiffness value was calculated for each specimen (Beer *et al.*). CT-based geometry of the internodal region was then extracted from the internodal region, and used to calculate the longitudinal Young’s modulus of the rind, E’ (Al-Zube *et al.*, 2018). The remaining material properties were calculated based on ratios derived from prior research, tests performed in our lab, and judicious estimates. The transverse Young’s modulus (E) was calculated as E’/100; the transverse shear modulus (G) was calculated as E’/1000, the longitudinal shear modulus (G’) was calculated as E’/10. These estimation rules were used only to obtain initial values for each parameter. As described in the following section, parametric analysis was used to independently vary these material properties for each specimen-specific *in silico* experiment. Prior sensitivity studies from our research group (Von Forell *et al.*, 2015; Stubbs *et al.*, 2019) showed that the longitudinal and transverse Poisson’s ratios (ν’, ν) of both the rind and the pith had no noticeable effects on model behavior. As a result, these parameters were not varied in this study.

### In Silico Sensitivity Analyses

*In silico* experiments produced an estimate of flexural stiffness and critical buckling failure load for each specimen-specific finite element model. Sensitivity analyses were performed by computing the relative change in these two outcomes resulting from small changes in material and geometric parameters. Each model was first analyzed with its own baseline material properties. Next, 8 additional analyses were performed. In each of these cases, one of the 8 material properties (E’, E, G’, and G for both rind and pith) was increased by 1%. Normalized sensitivity values were computed for each material property using the same method described in a previous study from our research group (Von Forell *et al.*, 2015).

Geometric sensitivities were obtained using the same equation, but using individual specimens that were chosen in pairs to exhibit negligible differences in two of the primary geometric parameters, but a significant difference (i.e. more than 7%) in the remaining parameters. This approach relied upon geometric data extracted from the 980 specimens of the previous study, and yielded one such pair for each of three geometric parameters, thus resulting in a sample size of 6 specimen-specific models. To control for potential material interaction effects, all of these models were assigned the same set of material properties.

For the linear buckling study, 12 specimen-specific finite element models were analyzed for 9 material variation cases. For the flexural stiffness study, 11 specimen-specific finite element models were analyzed for 9 material variation cases. As such, a total of 207 analyses were performed.

### Statistical Sensitivity Analyses

As the computational sensitivity analyses of the geometric parameters were based on a single comparison each (n = 1), statistical analyses of the specimen-specific *in vitro* tests were performed to verify the results. Statistical analyses were performed on the *in vitro* three-point bending data, which consisted of three predictor variables: major diameter, minor diameter, and rind thickness, as well as two response variables: flexural stiffness and maximum bending strength. Each variable was first non-dimensionalized by its respective mean value. Due to high variance inflation factors, univariate regression was performed between each predictor and response variable for a total of 6 univariate regressions (each of which had *n* = 956 data points).

## Results

### Validation

The response of *in silico* specimen-specific models closely matched the behavior of their physical counterparts. The correlation between physical and *in silico* values of flexural stiffness was very high (R^2^ = 0.99), and the flexural stiffness predictions were quantitatively accurate. The critical loads obtained from the *in silico* linear buckling experiments were compared to the critical loads of the *in vitro* three-point bending specimens, and found to have an R^2^ value of 0.56. These results are shown in Figure 2. As expected from a linear stability analysis, the predicted critical buckling failure loads were significantly higher than the actual loads at failure. This is because the linear buckling analysis does not account for factors such as cross-sectional ovalization and tissue failure which are influential in actual buckling, but that are not captured by linear buckling analysis. Finally, as will be explained in the following section, the computed sensitivities for geometric factors were similar for both physical and *in silico* approaches.

**Figure 2.**
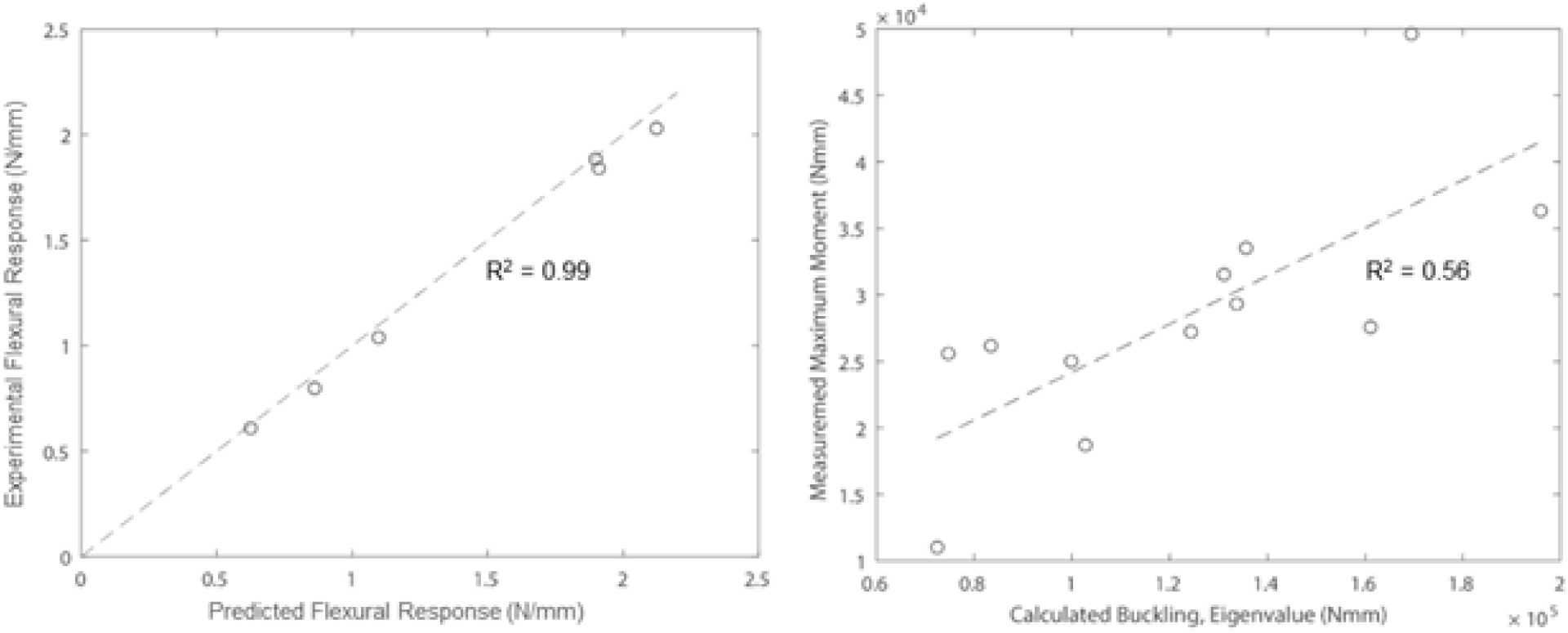
Model validation correlation plots, consisting of comparisons between predicted and measured values of flexural stiffness (left); and buckling load (right).

### In Silico Sensitivity Results

Sensitivity analyses of the *in silico* experiments were performed to determine the effect of each geometric and material parameter on the buckling strength and flexural stiffness of the specimens. The mean sensitivities for each parameter are shown in Figure 3. Geometric parameters were consistently found to be more influential than any of the material parameters for both analyses. The rind tissue properties generally had a larger impact on flexural stiffness than the pith material parameters. Of the geometric parameters, minor diameter was the most significant, major diameter was the second most significant, and rind thickness was the least significant.

**Figure 3.**
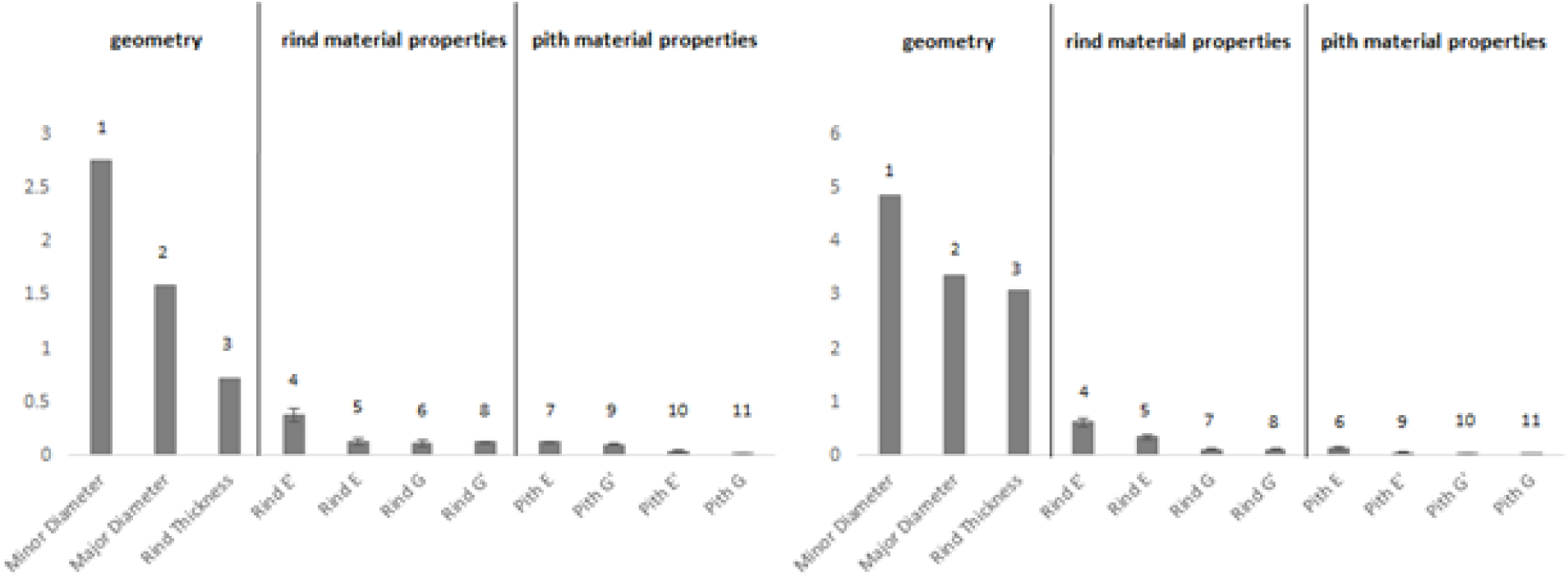
Normalized sensitivities (% change in output / % change in parameter) of geometric and material parameters for linear buckling (left) and flexural stiffness (right). The numeral above each bar indicates that parameter’s ordinal rank.

The mean sensitivities for each material parameter are shown in Figure 4, along with whiskers indicating the 95% confidence interval. The longitudinal Young’s modulus of the rind was the most significant material property for both types of flexural stiffness. The rank ordering of the material parameters differs slightly between the flexural stiffness and buckling studies.

**Figure 4.**
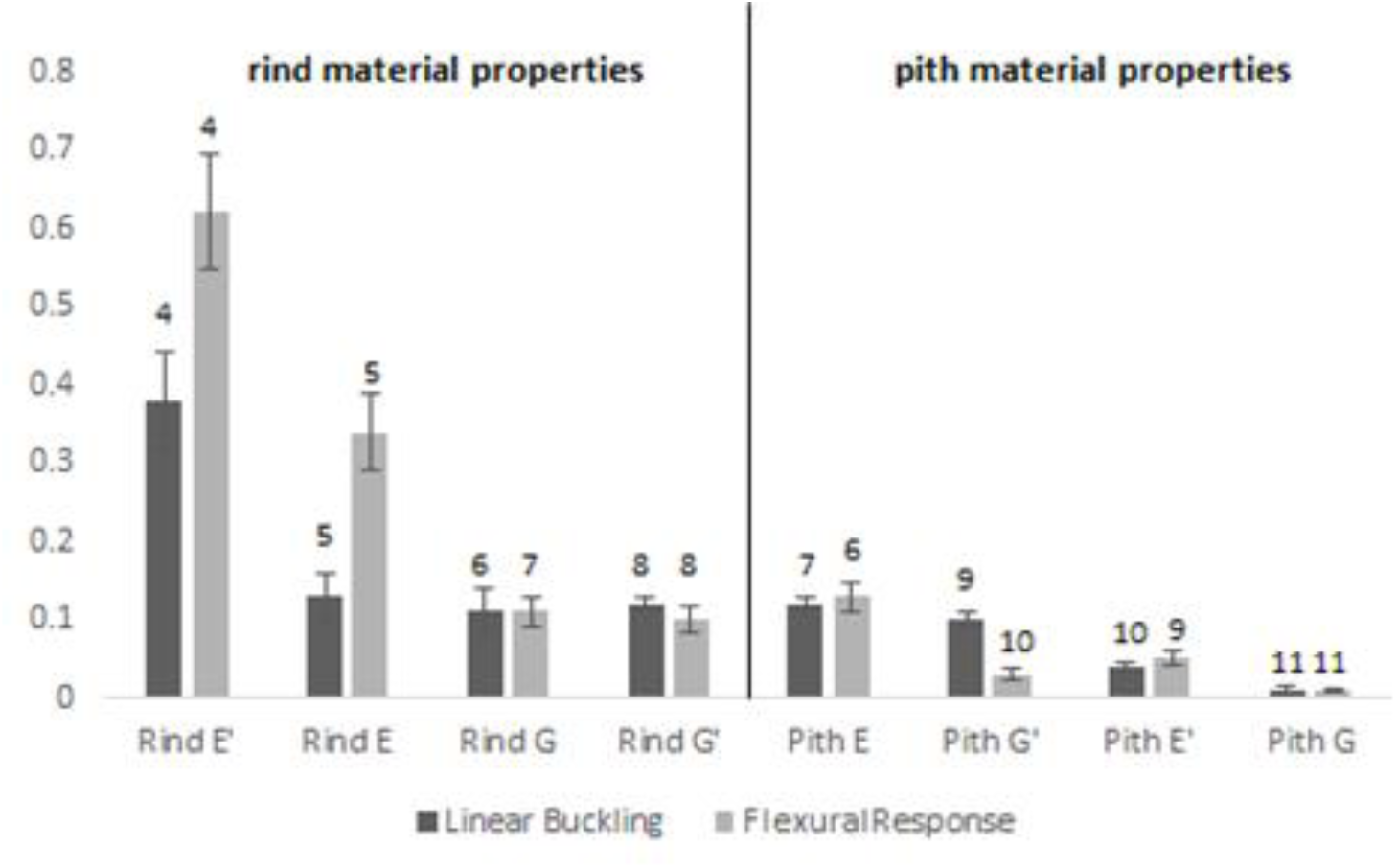
Normalized sensitivities (% change in output / % change in parameter) of material parameters for linear buckling and flexural stiffness, with 95% confidence intervals on the means

### Statistical Sensitivity Results

*In vitro* statistical sensitivities were assessed using the slope of univariate regressions between geometric parameters and actual flexural stiffness. Since all parameters were normalized, this approach produces unitless sensitivity values. The statistical sensitivity analysis was based on 956 *in vitro* tests. Major and minor diameters were found to be the most influential parameters for both flexural stiffness and buckling cases. Rind thickness was the least influential geometric parameter for both analyses. The mean sensitivities for this analysis are shown in Figure 5.

**Figure 5.**
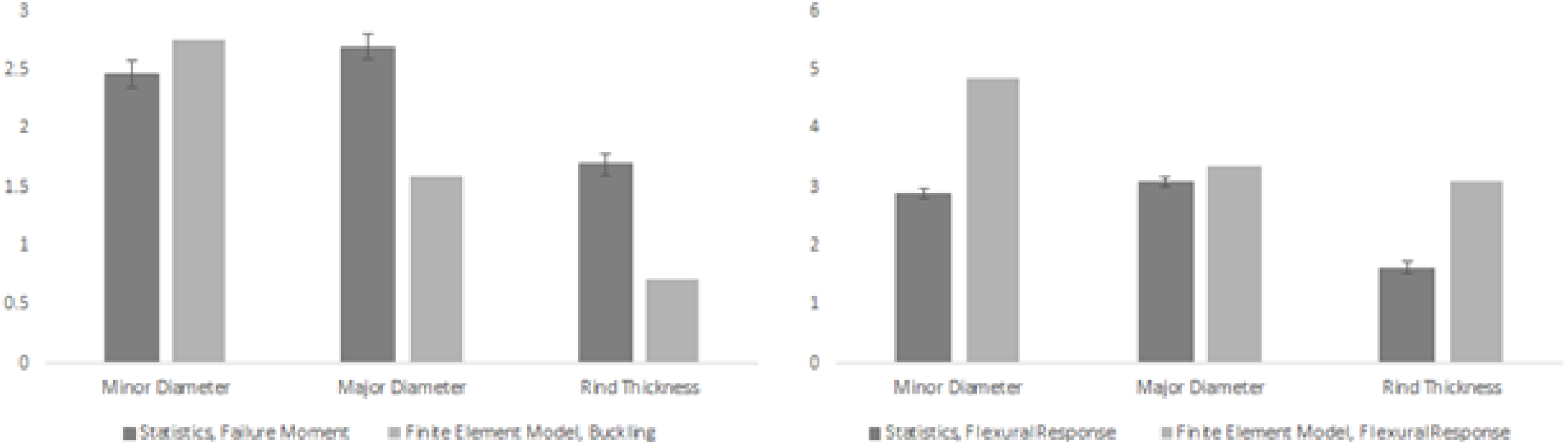
Normalized univariate regression (n = 956) coefficients of geometric parameters for linear buckling and flexural stiffness, with 95% confidence intervals on the regression coefficients, compared with finite element geometric sensitivities.

### Combined Results

All results can be combined into a single figure (Figure 6) that illustrates the sensitivity of maize stalk parameters to the parameters varied in this study. This figure clearly indicates the dominant effect of geometric parameters over those of the rind and pith.

**Figure 6.**
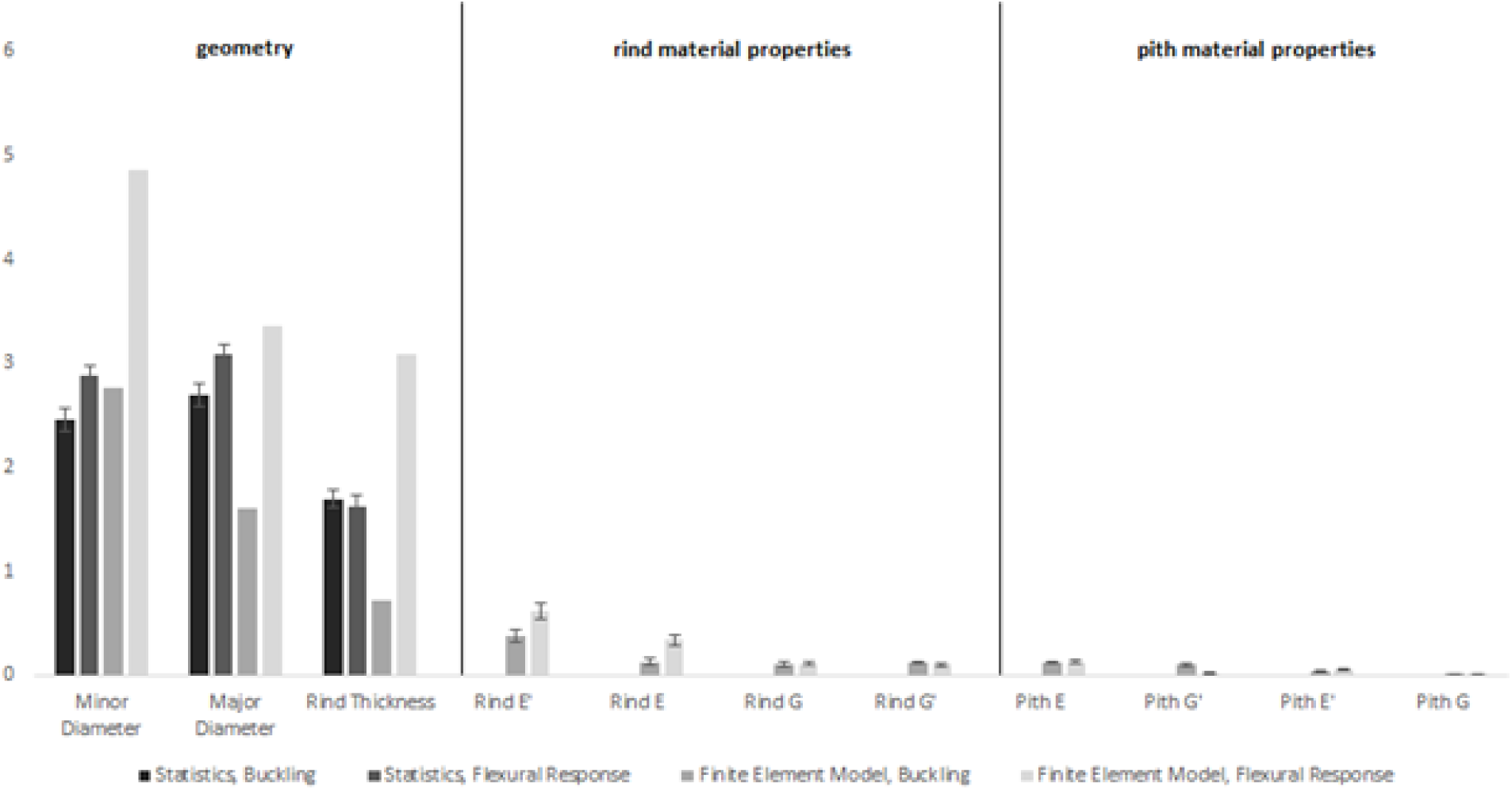
Combined plot of single-regression coefficients and normalized sensitivities of geometric and material parameters for linear buckling and flexural stiffness finite element model.

## Discussion

In the broader literature on maize lodging, chemical composition is often mentioned, but morphology and mechanical tissue properties are not commonly addressed. Yet from a structural engineering perspective, these factors are the most relevant for predicting most structural responses such as stiffness, strength, etc. This study demonstrates through both empirical and computational approaches that geometric factors have a strong influence on bending stiffness and bending strength of maize stalks, with material properties having a much lower influence.

A 2015 study from our research group obtained similar rankings as those presented above (Von Forell et al., 2015). However, the results in this study are more robust for several reasons. First, the models used in this study were thoroughly validated against experimental data (the models in the previous study were not). Second, this study used a broader sample of model geometries and used geometric factors that were more representative of the actual maize morphology (the previous study used a fairly limited geometric sampling). Thirdly, this study used a transversely isotropic material model for the pith tissue (as opposed to the isotropic model used in the prior study). Finally, and perhaps most importantly, the structural response of interest in (Von Forell *et al.*, 2015) was maximum bending stress, not the critical buckling load or flexural stiffness. But in spite of these differences, both studies identified geometric features as most influential and the minor diameter as the most influential geometric parameter. This provides further support for the generalization that the strength of maize stalks is primarily determined by morphological features, with mechanical tissue properties having a relatively weak influence on stalk strength.

However, more work will be required to understand the specific modes and mechanisms of failure in maize stems to better understand which continuum mechanics theories best predict the behavior and failure of the system. In particular, more work needs to be done to quantify how these relationships change with irregular morphologies and the details of how the pith influences stalk strength.

### Connections with structural engineering

To better understand the mechanisms behind maize stem failure, we can compare these results with the underlying continuum mechanics. For the simplified case of a hollow thin-walled cylinder in bending, the critical bending moment that induces tissue failure (M_t_) can be stated in terms of the fracture strain of the tissue (ε_F_’), longitudinal Young’s modulus of the material (E’), radius (r), and wall-thickness (t) by: *M*_*t*_ = *πϵ*_*F′*_*r*^2^*t* (Beer *et al.*). Similarly, the critical moment that induces buckling (M_b_) can be stated in terms of the radius (r), wall-thickness (t), longitudinal Young’s modulus (E’), and transverse Young’s modulus (E) by: 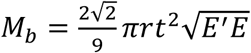 (Schulgasser and Witztum, 1992).

The sensitivity of each expression for critical load to each parameter can be obtained by differentiation and then normalized by multiplying by the parameter itself and divided by the expression for critical load. When this is carried out for the buckling expression, we see that geometric terms have normalized sensitivity values of 1 or greater while the two material parameters both have a normalized sensitivity of ½. Although maize stems are possess irregular cross-sectional geometry, and are not hollow, these analytical equations further support our conclusion that geometric features have more of an influence on the critical bending moment than material properties. These continuum mechanics theories seems to also be in agreement with previous studies which show that flexural stiffness, defined as the product of the longitudinal Young’s modulus (E’) and the second moment of area (I), is highly correlated with the overall strength of maize stalks (Robertson *et al.*, 2017).

### Limitations

This study relied upon geometrically realistic, but relatively simple models. The *in vitro* statistical models were univariate in nature and the *in silico* finite element models did not include non-linear effects such as large deformations or tissue failure. The material properties investigated in this study were all linear stiffnesses. This study did not investigate the influence of tissue *strength* on buckling as, at this time, there are no known studies that have measured this property.

At the same time, it should be restated that the geometric sensitivity analysis results were entirely empirical and based on an extensive set of tests. More advanced models and experiments will be necessary to provide further insights into issues such as the onset of buckling, and the role of tissue failure. But the trends shown above are unlikely to be affected in a major way.

In performing parametric sensitivity analyses, significant mechanistic insight into the system can be achieved. This method allows us to rank order each parameter by its individual effect on each complex phenotype or failure mode of the system, a task that is often impractical through experimentation of specimens with unknown material properties and complex geometries. It should be noted, however, that this approach is limited by the fact that these hypotheses drive towards a mechanistic understanding of each parameter individually. Although helpful, this ignores complex relationships that may between each parameters. It is undoubtable that the material properties and geometry of a maize stem are bonded together, being influenced by some of the same biomass allocation trade-offs, biotic influences, and abiotic factors throughout the plant’s lifespan, and by the interactions of those factors with the constantly-changing plant itself. Ultimately, a marriage of the individual influences of the parameters with the interrelationship between those parameters and their environmental context is necessary in building a complete understanding of the system.

## Conclusion

Stalk morphology has a much greater influence on the structural robustness of maize stalks than material stiffnesses. The major and minor diameters were found to be the most influential parameters, followed by the rind thickness. Of the material properties examined in this study, the longitudinal Young’s modulus of the rind was found to be the most influential, but material properties generally exhibited a relatively weak influence on structural response. The dominance of geometric parameters in the determination of the structural robustness of maize stems was supported in this study by experimental, computational, and theoretical analyses. This conclusion in consistent with other studies in the literature, though most studies indirectly address the difference in sensitivity between these parameter categories. (Schulgasser and Witztum, 1992; Wegst and Ashby, 2007; Stubbs *et al.*, 2018; (Maranville and Clegg; Von Forell *et al.*, 2015; Robertson *et al.*, 2016, 2017; Stubbs *et al.*, 2018).

This study provides clear evidence in support of the idea that deficiencies resulting from targeted reductions in organic polymers (such as lignin) may be counterbalanced by adjusting stalk morphology. Drawing on state-of-the-art techniques from two disciplines, plant scientists and engineers could work together to *design* a crop stalk architecture that would enable first/second generation biofuel production: crops that are high yielding, structurally robust, and whose biomass is readily converted to biofuel.

## Author Contributions

All authors were fully involved in the study and preparation of the manuscript. The material within has not been and will not be submitted for publication elsewhere.

## Acknowledgements

We thank Monsanto Company, St. Louis, MO, USA for providing the maize stalk samples used in this study. This work was funded in part by the National Science Foundation (Award # 1400973).

## References

Al-Zube L, Sun W, Robertson D, Cook D. 2018. The elastic modulus for maize stems. Plant Methods 14, 11.

Badel E, Ewers FW, Cochard H, Telewski FW. 2015. Acclimation of mechanical and hydraulic functions in trees: impact of the thigmomorphogenetic process. Frontiers in Plant Science 6.

Beer FP, Russell Johnston E, DeWolf JT, Mazurek DF. Mechanics of materials. 6th ed. New York: Mc Graw Hill; 2012.

Hibbitt K, Karlsson BI, Sorenson EP. 2016. ABAQUS/Standard theory manual. Sorenson Inc.

Hufner DR, Accorsi ML. 2009. A progressive failure theory for woven polymer-based composites subjected to dynamic loading. Composite Structures 89, 177–185.

Júnior MV, de Souza Neto EA, Munoz-Rojas PA. 2011. Advanced computational materials modeling: from classical to multi-scale techniques. John Wiley & Sons.

Maranville J, Clegg M. Morphological and physiological factors associated with stalk strength. In: G. Rosenberg, editor, Sorghum root and stalk rots: A critical review. Proceedings of the Consultative Group Discussion on Research Needs and Strategies for Control of Sorghum Root and Stalk Rot Diseases. Bellagio, Italy: ICRISAT, Patancheru, India; 1984. p. 111–118.

Matthews FL, Davies GAO, Hitchings D, Soutis C. 2000. Finite element modelling of composite materials and structures. Elsevier.

Moulia B, Coutand C, Julien J-L. 2015. Mechanosensitive control of plant growth: bearing the load, sensing, transducing, and responding. Frontiers in Plant Science 6.

Niklas KJ, Spatz H-C. 2012. Plant physics. University of Chicago Press.

Pedersen JF, Vogel KP, Funnell DL. 2005. Impact of reduced lignin on plant fitness. Crop Science 45, 812–819.

Robertson DJ, Julias M, Gardunia BW, Barten T, Cook DD. 2015a. Corn Stalk Lodging: A Forensic Engineering Approach Provides Insights into Failure Patterns and Mechanisms. Crop Science 55, 2833–2841.

Robertson DJ, Julias M, Lee SY, Cook DD. 2017. Maize Stalk Lodging: Morphological Determinants of Stalk Strength. Crop Science 57, 926–934.

Robertson DJ, Lee SY, Julias M, Cook DD. 2016. Maize Stalk Lodging: Flexural Stiffness Predicts Strength. Crop Science 56, 1711–1718.

Robertson DJ, Smith SL, Cook DD. 2015b. On Measuring the Bending Strength of Septate Grass Stems. American Journal of Botany 102, 5–11.

Sandorff PE. 1980. Saint-Venant effects in an orthotropic beam. Journal of Composite Materials 14, 199–212.

Schulgasser K, Witztum A. 1992. On the strength, stiffness and stability of tubular plant stems and leaves. Journal of theoretical biology 155, 497–515.

Simmons BA, Loqué D, Ralph J. 2010. Advances in modifying lignin for enhanced biofuel production. Current opinion in plant biology 13, 312–319.

Simulia DS. 2016. ABAQUS Analysis Manual. Providence, RI.

Stubbs CJ, Baban NS, Robertson DJ, Alzube L, Cook DD. 2018. Bending stress in plant stems: models and assumptions. Plant Biomechanics. Springer, 49–77.

Stubbs CJ, Sun W, Cook DD. 2019. Measuring the transverse Young’s modulus of maize rind and pith tissues. Journal of biomechanics 84, 113–120.

Von Forell G, Robertson D, Lee SY, Cook DD. 2015. Preventing lodging in bioenergy crops: a biomechanical analysis of maize stalks suggests a new approach. Journal of Experimental Botany 66, 4367–4371.

Wegst U, Ashby M. 2007. The structural efficiency of orthotropic stalks, stems and tubes. Journal of Materials Science 42, 9005–9014.

